# Structure Basis of Ca_v_1.1 Modulation by Dihydropyridine Compounds

**DOI:** 10.1101/2020.08.13.250340

**Authors:** Shuai Gao, Nieng Yan

## Abstract

1,4-Dihydropyridines (DHP), the most commonly used antihypertensives, function by inhibiting the L-type voltage-gated Ca^2+^ (Ca_v_) channels. DHP compounds exhibit chirality-specific antagonistic or agonistic effects. Recent structural elucidation of rabbit Ca_v_1.1 bound to an achiral drug nifedipine reveals the general binding mode for DHP drugs, but the molecular basis for chiral specificity remains elusive. Here, we report five cryo-EM structures of nanodisc-embedded Ca_v_1.1 in the presence of the bestselling drug amlodipine, a DHP antagonist (R)-(+)-Bay K8644, and a titration of its agonistic enantiomer (S)-(-)-Bay K8644 at resolutions of 2.9-3.4 Å. The amlodipine-bound structure reveals the molecular basis for the high efficacy of the drug. All structures with the addition of the Bay K8644 enantiomers exhibit similar inactivated conformations, suggesting that the agonistic effect of (S)-(-)-Bay K8644 might be transient. The similarity of these structures to that obtained in detergent micelles alleviates the concerns about potential structural perturbation by detergents.

## Introduction

Voltage-gated Ca^2+^ (Ca_v_) channels (VGCC) are responsible for a broad spectrum of physiological events, including muscle contraction, secretion, and synaptic signal transduction (*1, 2*). In mammals, 10 subtypes of Ca_v_ channels are classified to three subfamilies: Ca_v_1 (Ca_v_1.1-Ca_v_1.4), Ca_v_2 (Ca_v_2.1-Ca_v_2.3), and Ca_v_3 (Ca_v_3.1-Ca_v_3.3). Ca_v_1 channels, also known as the L-type VGCCs or dihydropyridine (DHP) receptors (DHPRs), are composed of a core α1 subunit and three auxiliary subunits, α2δ, β, and γ (*3, 4*). The α1 subunit is a single polypeptide of ∼ 2000 amino acids, folding into four homologous repeats I-IV. Each repeat contains six transmembrane segments (S1-S6) that form two functional entities: segments S1-S4 in each repeat constitute the peripheral voltage sensing domains (VSDs), and the S5-S6 segments from all four repeats, together with the intervening pore helices P1 and P2, enclose the central ion-permeating pore domain (PD). The short fragments between P1 and P2 from the four repeats serve as the molecular sieve, known as the selectivity filter (SF), which discriminates calcium from other ions (*1-4*).

Dysfunctional Ca_v_ channels are associated with various pathophysiological conditions ranging from cardiovascular disorders to psychiatric and neurological syndromes, such as cardiac arrhythmias, seizures, epilepsy, autism and Parkinson’s disease (*1-4*). Antagonists of DHPRs have demonstrated excellent efficacy in clinical practice for the treatment of specific conditions, including hypertension, cardiac ischemia, pain and tremor (*5*).

Among the Ca_v_ antagonists, DHP compounds are the most widely prescribed drugs, which, exemplified by amlodipine and nifedipine, have been the world’s bestsellers for decades (*6, 7*). Benzothiazepines (BTZ) and phenylalkylamines (PAA) represent the other two major classes of DHPR-targeting drugs (*6, 8, 9*). While BTZ and PAA compounds directly block ion conduction by traversing the central cavity of the pore domain, DHP antagonists bind to the fenestration, which is the portal on the side wall of the PD, on the interface of repeats III and IV for allosteric modulation (*9*).

While DHP drugs act as antagonists, some DHP compounds exhibit agonistic effects on DHPRs. Even more intriguingly, the mode of action (MOA) of some DHP compounds is stereo-selective; enantiomers of these DHP ligands possess opposite pharmacological profiles (*10*). For instance, whereas compound (*S*)-(-)-Bay K8644 is a potent agonist for DHPRs, its enantiomer, (*R*)-(+)-Bay K 8644 displays antagonistic activity (Fig. 1A) (*11, 12*). It is also noted that (*S*)-(-)-Bay K8644 produces a biphasic dose-response effect, whereby the agonist is turned to an antagonist when applied at high concentrations (*13, 14*). The biphasic property is also observed for modulators of other membrane proteins, such as dopamine D-2 and cannabinoid receptors, for which the usage dose should be finely regulated to achieve the desired efficacy (*15, 16*).

**Fig. 1.**
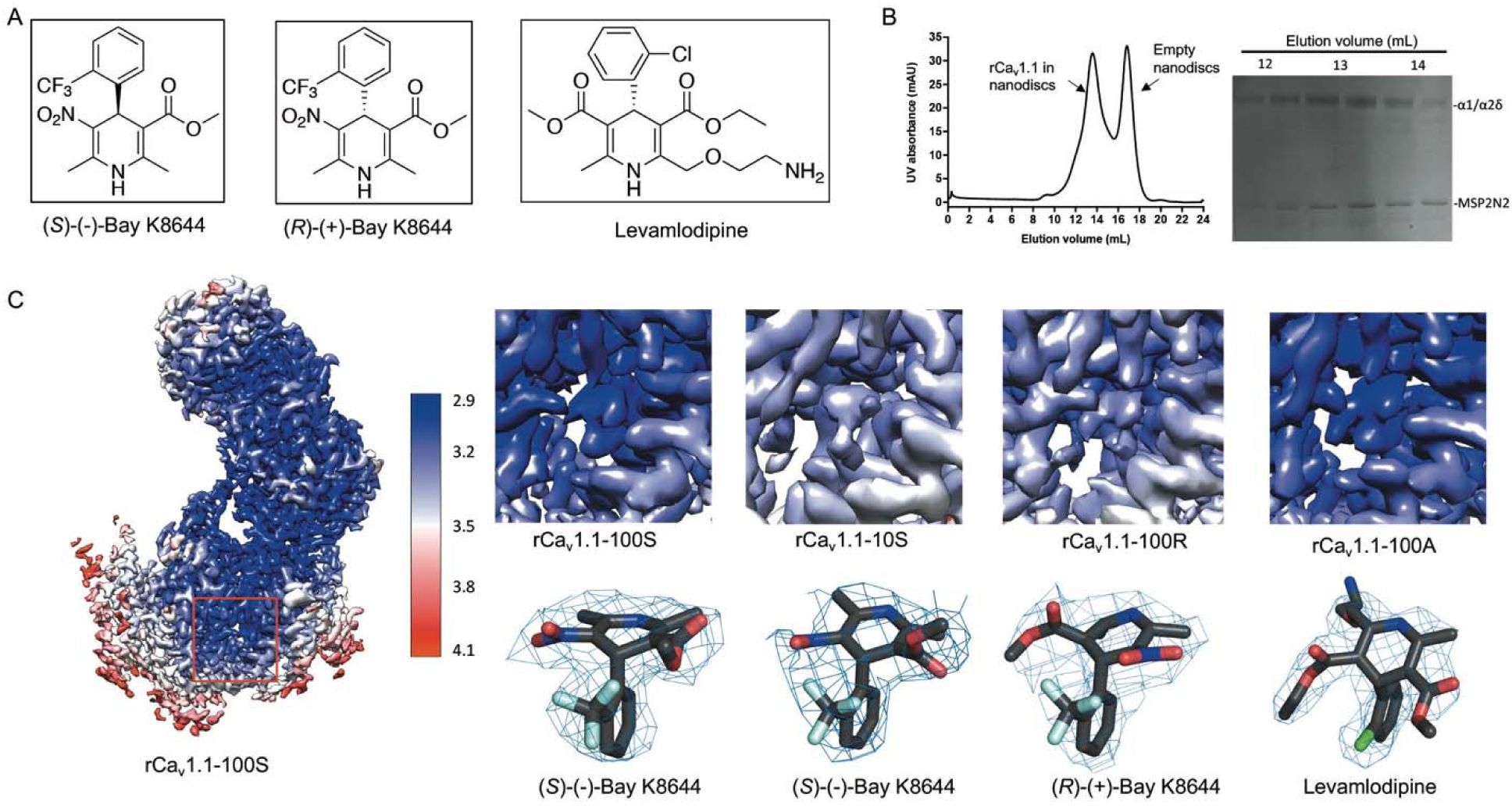
Cryo-EM analysis of Ca_v_1.1 in complex with amlodipine, RBK, and SBK in nanodiscs. (**A**) Chemical structures of (*S*)-(-)-Bay K8644, (*R*)-(+)-Bay K8644, and levamlodipine. (**B**) Final purification step for rCa_v_1.1 in MSP2N2-surrounded nanodiscs. Shown here is a representative size-exclusion chromatogram (SEC) and Coomassie blue-stained SDS-PAGE. (**C**) Local resolution maps and densities for the bound ligands. The densities in the lower row, shown as blue mesh, are contoured at 3 σ and prepared in PyMol.

An in-depth understanding of the MOA of these modulators requires high-resolution structures. Due to the advances of the resolution revolution of single-particle cryogenic electron microscopy (cryo-EM), we were able to resolve the structures of the rabbit Ca_v_1.1 (rCa_v_1.1) channel complex and human Ca_v_3.1 alone and in the presence of various small molecule ligands (*17-20*). Specifically, the atomic structures of rCa_v_1.1 bound to representative DHP, BTZ, and PAA drugs, nifedipine, diltiazem, and verapamil, respectively, have elucidated the molecular details of the drugs’ action (*19*).

Notwithstanding these exciting structural advances, there are a number of outstanding questions. First, all the reported structures of eukaryotic Ca_v_ channels and the closely related Na_v_ channels are of proteins purified in detergent micelles. It is unclear whether the detergents have altered local structures of these highly dynamic channels. As such, structural elucidation in a more physiologically relevant environment, such as in nanodiscs, is required. Second, in the structure of Ca_v_1.1 complexed with the agonist (*S*)-(-)-Bay K8644, the overall structure conforms to what is expected to be an inactivated state (*19*). (*S*)-(-)-Bay K8644 was applied at 100 μM, a dose at which the compound may function as an antagonist. It remains to be tested whether the agonist at lower concentration can help lock the channel in an activated state. Last but not least, both DHP compounds, (*S*)-(-)-Bay K8644 and nifedipine, have relatively small chemical groups, but it has yet to be seen how the bulkier groups of some DHP drugs, such as levamlodipine (amlodipine for short hereafter), are coordinated by DHPR.

To address these remaining questions, we sought to resolve the structures of rabbit Ca_v_1.1 reconstituted in nanodiscs with addition of representative DHPR compounds. Here we report high-resolution cryo-EM structures of nanodisc-embedded Ca_v_1.1 bound to the antagonists amlodipine and (*R)*-(+)-Bay K 8644, and a titration of (*S*)-(-)-Bay K8644 (Fig. 1A). These structures together provide advanced knowledge on the modulation of DHPRs by DHP compounds. For simplicity, we will refer to the enantiomers of Bay K 8644 as RBK and SBK hereafter.

## Results

### Structures of Ca_v_1.1 in lipid nanodiscs are nearly identical to those in detergent micelles

Following our published protocols (*17, 21*), Ca_v_1.1 isolated from the skeletal muscle of New Zealand white rabbits, rCa_v_1.1, was purified and reconstituted into nanodiscs with the membrane scaffold protein 2N2 (MSP2N2) and POPC (Fig. 1B, S1). Please refer to Methods for details of the protein purification and nanodisc reconstitution. The mono-dispersed peak fractions of rCa_v_1.1 nanodiscs from size-exclusion chromatography were pooled and incubated with the target molecules before cryo-sample preparation. Amlodipine and RBK were applied at 100 μM, and SBK was added at final concentrations of 1 μM, 10 μM, and 100 μM, in the hope of capturing different channel states.

The cryo-grids were made following a standard protocol and electron micrographs were collected on Titan Krios G3 cryo-electron microscope equipped with the spherical aberration (Cs) image corrector and GIF quantum electron energy filter. The workflow for data processing is described in SI (Fig. S2, S3). The overall resolutions of the channel complexes were determined at 2.9 Å with amlodipine, 3.2 Å with RBK, and 3.4 Å, 3.4 Å, and 3.0 Å with SBK at 1 μM, 10 μM, and 100 μM, respectively. For simplicity, we will refer to these five structures as rCa_v_1.1-100A (with 100 μM amlodipine), 100R (with 100 μM RBK), and 1S/10S/100S (with SBK applied at three different concentrations). Our published structure of digitonin-embedded rCa_v_1.1 in the presence of 200 μM nifedipine, used as structural reference several times in this manuscript, will be referred to as rCa_v_1.1-200N (PDB code: 6JP5) (*19*). The excellent map quality and high local resolutions allow for accurate assignment of the DHP ligands (Fig. 1C, S4).

The overall structures of rCa_v_1.1 in complex with the DHP antagonists, amlodipine and RBK, in nanodiscs are nearly identical to rCa_v_1.1-200N. The structures of the α1 subunit in rCa_v_1.1-100A and rCa_v_1.1-100R can be superimposed to that in rCa_v_1.1-200N with root-mean-square deviation (RMSD) values both of 0.52 Å over 1115 Cα atoms (Fig. 2A, B). This observation in part supports that the previous structure-function relationship studies of rCa_v_1.1 obtained in a detergent surrounding can be repeated in lipid membrane environment.

**Fig. 2.**
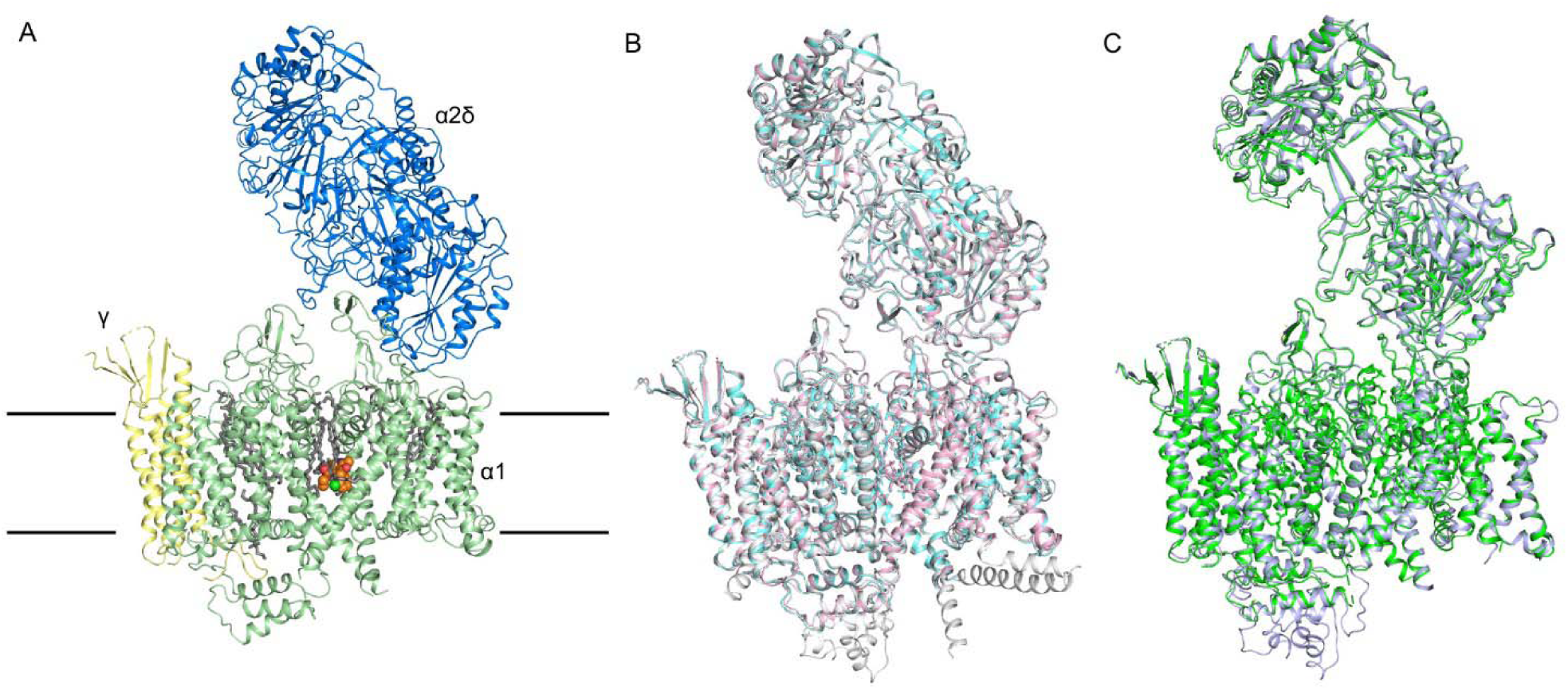
Nearly identical conformations of DHP antagonists-bound rCa_v_1.1 in detergents and in nanodiscs. (**A**) Overall structure of the amlodipine-bound rCa_v_1.1 complex (rCa_v_1.1-100A) at 2.9 Å resolution. The overall structure of the channel complex is shown for different subunits. The β1 subunit is omitted throughout the manuscript because of its poor resolution. Amlodipine is shown as orange spheres and the bound lipids are shown as grey sticks. (**B**) The overall structures of rCa_v_1.1 in complex with amlodipine (pink) and RBK (cyan) in nanodiscs and nifedipine (grey) in digitonin are nearly identical. (**C**) Structures of rCa_v_1.1 in complex with 100 μM SBK in nanodisc (green) and in digitonin (blue) are nearly identical.

The DHP agonist, SBK, was titrated into the purified rCa_v_1.1 embedded in nanodiscs at the final concentration of 1μM, 10 μM, and 100 μM, respectively, immediately before cryo-sample preparation. In all three nanodisc-surrounded reconstructions, the intracellular gate is closed and the four VSDs exhibit depolarized conformations, characteristic of the same putative inactivated state as our previously reported SBK-bound rCa_v_1.1 in digitonin (PDB code: 6JP8) (*19*). The overall structures of rCa_v_1.1-100S in nanodiscs and in detergents can be well aligned with a root-mean-square deviation (RMSD) values of 0.55 Å over 1115 Cα atoms for α1 subunit (Fig. 2C). This observation partially alleviates the concern with potential interference of the conformations of Ca_v_1.1, and probably all other single-chain Ca_v_ and Na_v_ channels whose structures have been resolved, by detergents.

### Nearly identical state of rCa_v_1.1 in complex with a titration of SBK

As aforementioned, the DHP agonist, SBK, was added into nanodisc-embedded rCa_v_1.1 at a concentration gradient with the aim to capture different channel states. The EM densities corresponding to SBK are only observed in rCa_v_1.1-10S/100S, leaving the rCa_v_1.1-1S as an apo reconstruction (Fig. 3A). SBK is nestled into the same fenestration site enclosed by the pore-forming elements of repeats III and IV, surrounded by residues on the segments P1_III_, S5_III_, S6_III_ and S6_IV_ (Fig. 3B). Because the detailed coordination of SBK is identical to our reported one (*19*), it will not be elaborated here.

**Fig. 3.**
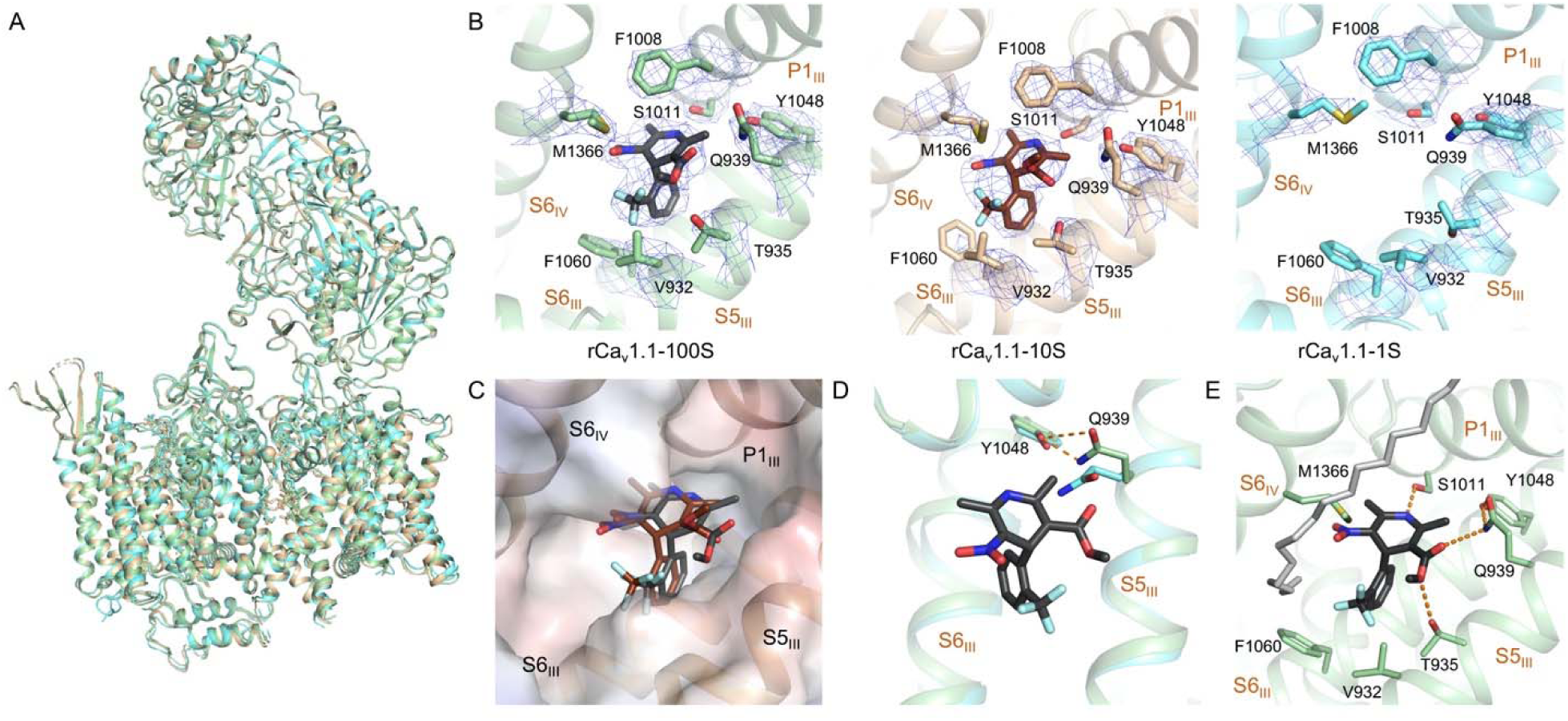
The conformation of nanodisc-embedded rCa_v_1.1 remains nearly unchanged with SBK applied at different conformations. (**A**) Structural comparison of rCa_v_1.1 in complex with 100 μM SBK (green), 10 μM SBK (wheat), and 1 μM SBK (cyan) in nanodisc. Despite the different concentrations of SBK applied, the overall structure remains nearly unchanged. (**B**) There is no density of SBK in the EM reconstruction when applied at 1 μM. The densities for SBK and its binding site in rCa_v_1.1-100S/10S/1S are shown. The maps, shown as blue mesh, are contoured at 7σ and prepared in PyMol. (**C**) Slight displacement of SBK between rCa_v_1.1-100S (black) and 10S (brown). The electrostatic surface potential of rCa_v_1.1-100S was calculated in PyMol. (**D**) Gln939 on S5_III_ of rCa_v_1.1-100S (green) shifts to Tyr1048 compared to rCa_v_1.1-1S (cyan). Such local conformational shift is important for accommodating the ligand. (**E**) A lipid may contribute to SBK coordination. SBK and lipid are shown as sticks and colored black and gray, respectively. SBK is coordinated by both polar and hydrophobic residues. One hydrophobic tail of a phospholipid blocks the III-IV fenestration, strengthening SBK binding. Potential hydrogen bonds are shown as dashed lines.

The EM density for SBK in rCa_v_1.1-10S is worse than that in rCa_v_1.1-100S, implying less stable coordination. A close examination shows a slight difference in the densities for the bound SBK in rCa_v_1.1-10S and -100S, especially those for the C3-ester group. Structural models were built for SBK in rCa_v_1.1-10S, which displaces slightly from that in rCa_v_1.1-100S. In particular, the C3-ester group appears to undergo a rotation (Fig. 3C). This minor difference may not change its function as an antagonist to rCa_v_1.1 in this particular conformation. Nevertheless, this observation is consistent with our previous analysis on the meta-stable association between rCa_v_1.1 and SBK.

Due to the lack of ligand density, rCa_v_1.1-1S may represent the state of an apo-structure, hence providing a good control to trace the conformational changes of rCa_v_1.1 upon SBK binding. The only difference occurs in Gln939 on S5_III_ (Fig. 3D). Upon SBK entry, Gln939 rotates toward Tyr1048 of S6_III_ and mediates the essential H-bond network between the ester group of SBK and Tyr1048. It was reported that single point mutations corresponding to Y1048F and Y1048A resulted in reduced affinity with DHP by ∼10- and ∼1,000-fold, respectively (*22*). Our structural comparison suggests that Tyr1048 may be required for DHP association through facilitating the shift of Gln939. Rotation of Gln939 driven by Tyr1048 paves the path for the entry of DHP ligands, followed by the stabilization from the hydrophobic and H-bond interactions with the surrounding residues in the fenestration site.

It is noted that a density corresponding to a phospholipid was well-resolved outside the fenestration (Figure 3E, Fig. S5A). One aliphatic tail of the lipid blocks the fenestration and interacts directly with SBK. This arrangement may prevent the ligand from exiting the binding site. The role of phospholipid in ligand binding to membrane proteins has rarely been discussed. Our structural finding provides a clue to understanding the sophisticated ligand binding within membrane.

### Comparison for DHP enantiomers

SBK and RBK are a pair of DHP enantiomers with a chiral center on the C4 atom of the dihydropyridine ring. In rCa_v_1.1-100R, RBK is positioned in the same fenestration site as that for SBK in rCa_v_1.1-10S/100S (Fig. 4A). Compared to SBK in rCa_v_1.1-100S, the CF_3_-substituted phenyl ring of RBK projects into the hydrophobic pocket formed by Thr935, Val932, and Phe1060, while the dihydropyridine flips over ∼180 degree (Fig. 4B). In this way, the nitro group of RBK is H-bonded to Tyr1048/Gln939 and Thr935, similar to the role of C3-ester in SBK.

**Fig. 4.**
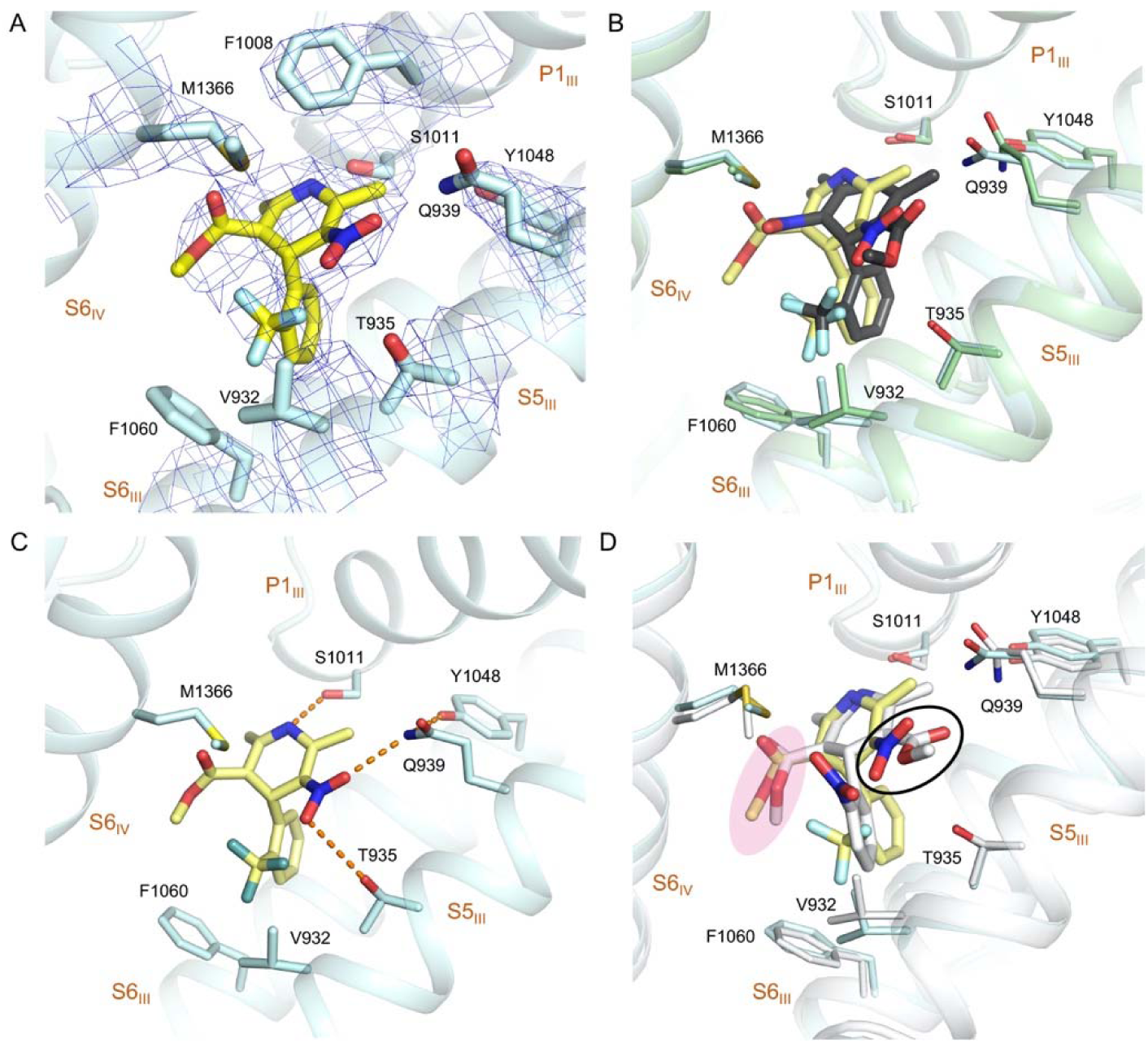
Coordination of the antagonistic RBK by rCa_v_1.1. (**A**) The densities for the RBK binding site in rCa_v_1.1-100R. RBK is shown as yellow sticks. The densities, shown as blue mesh, are contoured at 7 σ and prepared in PyMol. (**B**) Overlapping binding site for RBK and SBK. The structures of rCa_v_1.1-100R (blue) and -100S (green) are superimposed. RBK and SBK are colored yellow and black, respectively. (**C**) Coordination of RBK. Potential hydrogen bonds are shown as dashed lines. (**D**) Overlapping binding site for RBK and nifedipine. The structure rCa_v_1.1-100R (yellow) and nifedipine (gray) are superimposed. The NO_2_ group in RBK, which is positioned adjacent to S5_III_ segment, is highlighted with black circle. The C3-ester groups of RBK and nifedipine are highlighted with pink shadow.

On the other hand, the C3-ester of RBK is situated into a hydrophobic cavity surrounded by Met1366, Val932, and Phe1060 (Fig. 4C). The binding pose of RBK can be superimposed to that for nifedipine (PDB:5JP5), suggesting a conserved conformation for DHP antagonists (Fig. 4D). Besides, the cryo-EM coordination of RBK in nanodisc-embedded rCa_v_1.1 is consistent with our previous docking result using the aforementioned nifedipine-bound structure as template (*19*).

### Coordination of amlodipine

Amlodipine is the most potent DHP antagonist that displays a pH-dependent efficacy (*23*). As expected, it is also nestled in the III-IV fenestration of the PD. Despite the larger size, coordination for the backbone of amlodipine is nearly identical with that for nifedipine, involving residues Ser1011, Tyr1048, Gln939, Thr935, and Met1366 (Fig. 5A, B).

**Fig. 5.**
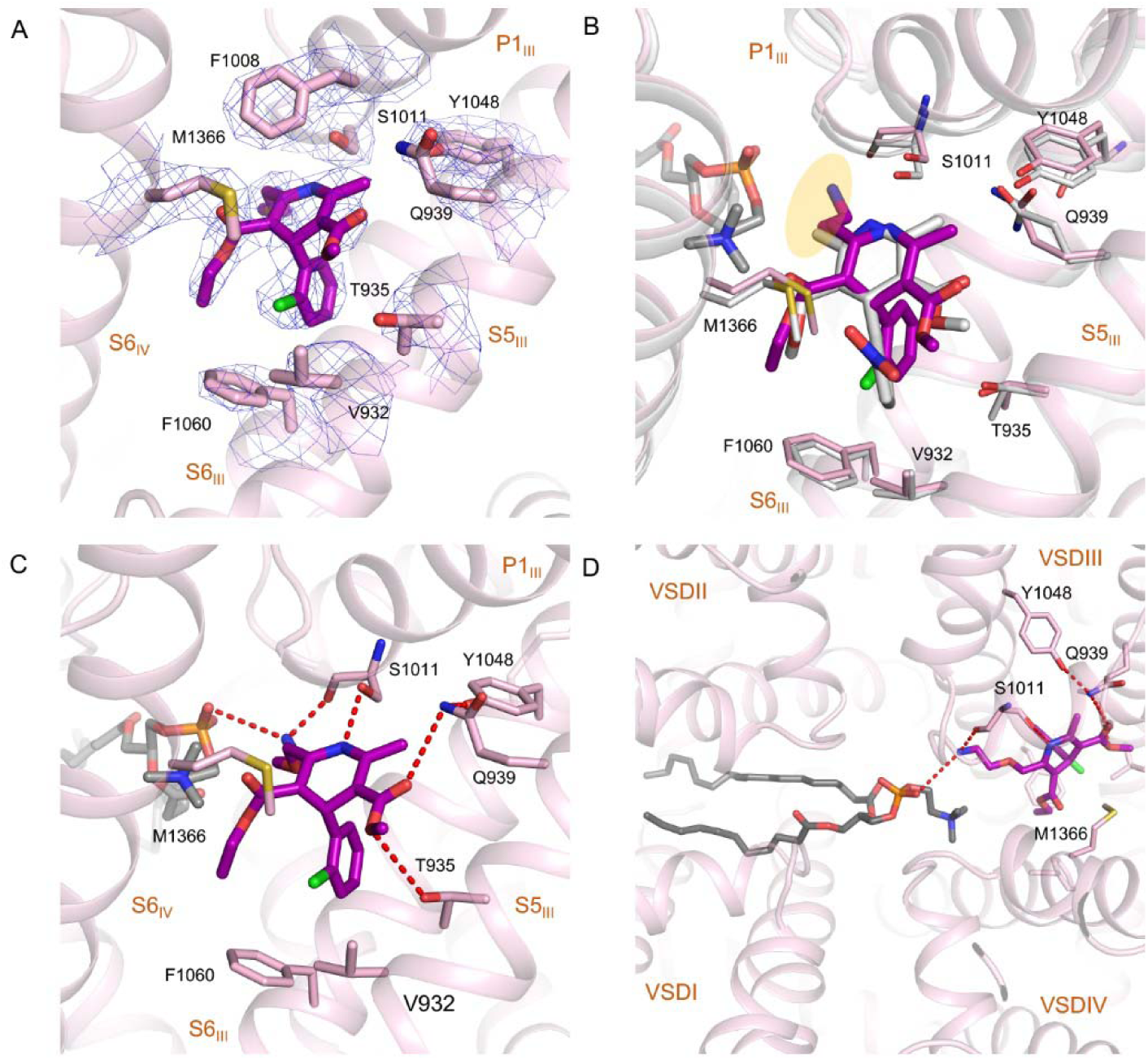
Specific interactions between rCa_v_1.1 and Amlodipine. (**A**) The density for amlodipine binding sites in rCa_v_1.1-100A. Amlodipine is shown as purple sticks. The densities, shown as blue mesh, are contoured at 7σ and prepared in PyMol. (**B**) Overlapping binding site for amlodipine and nifedipine. The structure rCa_v_1.1-100A (pink) and nifedipine (gray) are superimposed. The ethanolamine group in amlodipine is highlighted with orange shadow. (**C**) Coordination of amlodipine at the III-IV fenestration of rCa_v_1.1. (**D**) A transverse lipid in the central cavity contributes to amlodipine binding. The phosphate group (orange colored) of the lipid (grey for the backbone) and Ser1011 coordinate the ethanolamine group of amlodipine from two opposite directions. Please refer to figure S5B for the density of the lipid.

The bulky substituent in amlodipine, the ethanolamine group, points to the central cavity of the PD and is H-bonded to the carbonyl group of S1011 (Fig. 5A, B). Except for this group, the interactions between amlodipine and the surrounding residues are highly conserved similar to other antagonists in rCa_v_1.1, including three pairs of H-bonds, two between C3-ester and Tyr1048/Gln939/Thr935, and one between N1 atom and S1011.

Amlodipine displays longer duration time of action than nifedipine, allowing a once-a-day dosage regimen in human (*24*). Based on our structure, the improved pharmacological profile of amlodipine may come from the ethanolamine group. The introduction of ethanolamine group not only increases the aqueous solubility, but also enhances the binding affinity of amlodipine by the formation of an additional H-bond with the conserved residue in Ca_v_1 that corresponds to Ser1011 in rCa_v_1.1 (Fig. 5C).

### Potential role of lipids in DHP ligand coordination

In addition to the compound and surrounding residues, EM densities corresponding to lipids were resolved in the central cavity of DHP-bound rCa_v_1.1. It is noteworthy that such densities were also observed in the digitonin solubilized conditions for both ligand-free and antagonists-bound rCa_v_1.1 reconstructions. Although the resolution is insufficient to identify specific lipids, the contour allowed for putative docking of the lipid 1-palmitoyl-2-oleoyl-sn-glycero-3-phosphoethanolamine (16:0-18:1 PE), a major lipid component of eukaryotic cell membrane (*18*).

Given the proximity to the molecules, the lipid appears to directly contribute to the binding of amlodipine. Specially, in rCa_v_1.1-100A, the phosphate group of the lipid approaches the fenestration site and interacts with the ethanolamine substituent of amlodipine (Fig. 5C,D, Fig. S5B). Although the exact identity of this specific lipid cannot be identified due to technical limitations, the phosphate is the common functional group in all phospholipids, which indicates that the phosphate group of the native lipid may provide further stabilization for amlodipine (Fig. 5D).

## Discussion

rCa_v_1.1 was the first single-chain VGIC (voltage-gated ion channels) whose structure was determined (*17, 18*). We have been employing it as a prototype for cryo-EM analysis of VGIC members. Before this study, all the structures of rCa_v_1.1, as well as the closely related human Ca_v_3.1 and multiple eukaryotic Na_v_ channels, were solved as proteins embedded in detergents (*25-30*). We have had some concerns with potential structural perturbation of these highly dynamic molecular machines by detergents. The structural similarity of rCa_v_1.1 in nanodiscs and in detergent micelles, which was also observed in our recently published NaChBac (*31*), alleviates this concern and consolidates the structural findings obtained from the detergent-embedded channels. More satisfyingly, the coordination of the backbones of all resolved DHP antagonists, nifedipine (*19*), amlodipine, and RBK, is conserved regardless of the surrounding milieu of detergents or nanodiscs.

The high resolutions of rCa_v_1.1 in complex with different DHP compounds allowed for detailed analyses of ligand binding, providing the basis to understand the distinct efficacy of the drugs and functional groups of the channel. The direct participation of a phospholipid in the coordination of amlodipine is intriguing in that it is unclear whether a phospholipid can enter the central cavity in the intact membrane. Although our structure was obtained in nanodiscs, the native membrane was disrupted during protein extraction and purification. Therefore, this remains to be an enigma that awaits further investigation. Multiple biophysical and computational approaches are needed to dissect the mechanism and the function of the transverse lipids in the physiology and pharmacology of VGIC channels.

We had aimed to utilize the well-characterized agonist SBK to capture the channel in an activated state. Despite our attempt to apply SBK at different doses and in different environments, the channel remains in the same inactivated state. Notably, when applied at 1 μM, the density corresponding to SBK was gone. This observation supports our previous analysis that the SBK is not favored by the inactivated conformation. Therefore, a high concentration is required to compensate for the penalty (*19*). This set of structural analyses suggest that an agonist, whose action is to prolong the opening duration of the channel(*32-34*), is insufficient to lock the channel in an active conformation. Because it is impractical to conveniently introduce point mutations to the endogenous channels, it may require the invention of new methods to capture voltage-gated ion channels in distinct functional states.

Despite the remaining questions, our structural analyses reported here and previously (*19*) provide a more comprehensive understanding of the MOA of DHP drugs. This updated knowledge will facilitate drug discovery targeting other Ca_v_ channels and additional VGICs in general.

## Materials and Methods

### Expression and purification of GST-β1a

The GST-fused β1a protein was expressed and purified as previously reported (*17*). Briefly, overexpression of GST-β1a was carried out using *Escherichia coli* BL21 (DE3) (Novagen) transformed with plasmid containing the cDNA inserted into pGEX-4T-2 vector. The resulting *E*.*coli* was cultured at 37 °C in LB medium with 100 mg/mL ampicillin (AMRESCO) to OD_600_ of 1.2. Then the culture temperature was cooled down to 18 °C, and 0.2 mM isopropyl-β-D-thiogalactoside was added for protein expression induction. After 12 h culture, the cells were collected and resuspended with lysis buffer containing 25 mM Tris-HCl, pH 8.0 and 150 mM NaCl. Sonication was adopted for cell disruption. After centrifugation at 27,000 g for 1 h, the supernatant was collected and applied to Glutathione Sepharose 4B resin (GS4B, GE Healthcare) and washed with lysis buffer. The protein was eluted with elution buffer containing 50 mM Tris-HCl, pH 8.0, 50 mM NaCl and 10 mM glutathione. The ion-exchange chromatography was performed for further protein purification with an anion-exchange column (Source 15Q, GE Healthcare). Then peak fractions were collected and stored at -80 °C before using.

### Isolation of skeletal muscle membrane from New Zealand white rabbits

This step was well described previously (*17*). Fresh skeletal muscle was isolated from rabbit legs and back, and then was homogenized in buffer containing 0.3 M sucrose, 10 mM MOPS-Na, pH 7.4, 0.5 mM EDTA and protease inhibitors mixture including 2 mM phenylmethylsulphonyl fluoride (PMSF), 1.3 mg/mL aprotinin, 0.7 mg/mL pepstatin and 5 mg/mL leupeptin. Two steps centrifugation were performed. The homogenates were first centrifuged at 5,000 g for 5 min, following by a second centrifugation at 200,000 g for 1 h of the supernatant. The pellet was collected and stored at -80 °C before using.

### Protein preparation

The Ca_v_1.1 complex from rabbit skeletal muscle was purified as described previously. In brief, the collected skeletal muscle was solubilized in buffer containing 20 mM MOPS-Na, pH 7.4, 500 mM NaCl, 0.5 mM CaCl_2_, 1% (w/v) GDN (Anatrace), 2 mM PMSF, 3.9 mg/mL aprotinin, 2.1 mg/mL pepstatin, 15 mg/mL leupeptin and excessive GST-β1a at 4 °C for 2 h. The mixture was then ultra-centrifuged at 200,000 g for 30 min, and the supernatant was applied to GS4B resin. The protein-loaded resin was washed with buffer containing 20 mM MOPS-Na, pH 7.4, 500 mM NaCl, 0.5 mM CaCl_2_, 0.01% GDN and protease inhibitors. Protein elution was performed with buffer containing 100 mM Tris-HCl, pH 8.0, 200 mM NaCl, 0.5 mM CaCl_2_, 15 mM reduced glutathione, 0.01% GDN, and protease inhibitors. The fused GST was removed with addition of HRV 3C protease, followed by the size-exclusion chromatography (Superose 6 10/300 GL, GE Healthcare) in buffer containing 20 mM MOPS-Na, pH 7.4, 200 mM NaCl, 0.5 mM CaCl_2_, 0.01% GDN and protease inhibitors.

### Nanodisc reconstitution

Lipid 1-palmitoyl-2-oleoyl-glycero-3-phosphocholine (POPC, Avanti) in chloroform was dried under a nitrogen stream and resuspended in reconstitution buffer containing 25 mM Tris-HCl, pH 8.0, 150 mM NaCl, and 0.7% DDM. Approximately 250 μg purified protein was mixed with 1 mg MSP2N2 and 750 μg lipid. The mixture was incubated at 4 °C for 5 h with gentle rotation. Bio-beads (0.3 g) were then added to remove detergents from the system and facilitate nanodisc formation. After incubation at 4 °C overnight, Bio-beads were removed through filtration, and protein-containing nanodiscs were collected for cryo-EM analysis after final purification by SEC in running buffer containing 25 mM Tris-HCl, pH 8.0, and 150 mM NaCl (Superose 6 Increase 10/300 GL, GE Healthcare).

### Protein-drug sample preparation

For rCa_v_1.1-1S reconstruction, (*S*)-(-)-Bay K8644 was added into the nanodisc reconstitution buffer at the final concentration of 1 μM. The mixture was purified by SEC in running buffer supplemented with 1 μM (*S*)-(-)-Bay K8644.

For rCa_v_1.1-10S/100S/100R/100A reconstructions, 10 μM, 100 μM (*S*)-(-)-Bay K8644, 100 μM (*R*)-(+)-Bay K8644 and 100 μM amlodipine was separately incubated with nanodisc-embedded proteins for 30 min before cryo-grids preparation.

### Cryo-EM data acquisition

Aliquots of 3.5 μl concentrated samples were loaded onto glow-discharged holey carbon grids (Quantifoil Cu R1.2/1.3, 300 mesh). Grids were blotted for 5 s and plunge-frozen in liquid ethane cooled by liquid nitrogen using a Vitrobot MarK IV (Thermo Fisher) at 8 °C with 100 percent humidity. Grids were transferred to a Titan Krios electron microscope (Thermo Fisher) operating at 300 kV and equipped with a Gatan Gif Quantum energy filter (slit width 20 eV) and spherical aberration (Cs) image corrector. Micrographs were recorded using a K2 Summit counting camera (Gatan Company) in super-resolution mode with a nominal magnification of 105,000x, resulting in a calibrated pixel size of 0.557 Å. Each stack of 32 frames was exposed for 5.6 s, with an exposure time of 0.175 s per frame. The total dose for each stack was ∼ 50 e-/Å2. SerialEM was used for fully automated data collection (*35*). All 32 frames in each stack were aligned, summed, and dose weighted using MotionCorr2 and 2-fold binned to a pixel size of 1.114 Å/pixel (*36-38*). The defocus values were set from -1.9 to -2.1 μm and were estimated by Gctf (*39*).

### Image processing

Totals of 1739/2327/1595/1767/2092 cryo-EM micrographs were collected, and 644,294/714,871/560,063/621,407/686,403 particles were auto-picked by RELION-3.0 for rCa_v_1.1-100S, rCa_v_1.1-10S, rCa_v_1.1-1S, rCa_v_1.1-100R and rCa_v_1.1-100A in nanodiscs, respectively (*40*). Particle picking was performed by RELION-3.0. All subsequent 2D and 3D classifications and refinements were performed using RELION-3.0. Reference-free 2D classification using RELION-3.0 was performed to remove ice spots, contaminants, and aggregates, yielding 581,802/701,745/463,685/537,339/506,092 particles, respectively. The particles were processed with a global search K=1 using RELION-3.0 to determine the initial orientation alignment parameters using bin2 particles. A published cryo-EM map of human rCa_v_1.1 was used as the initial reference. After 40 iterations, the datasets from the last 6 iterations were subject to local search 3D-classifications using 4 classes. Particles from good classes were then combined and re-extracted with a box size of 280 and binned pixel size of 1.114 Å for further refinement and 3D-classification, resulting in 157,294/207,038/135,210/146,737/184,373 particles, which were subjected to auto-refinement. 3D reconstructions were obtained at resolutions of 3.1 Å/3.5 Å/3.8 Å/3.4 Å/3.1 Å. Skip-alignment 3D classification using bin1 particles after Bayesian polish yielded data sets containing 87,516/53,789/53,341/94,696/184,373 particles respectively and resulted in respective reconstructions at 3.0 Å/3.4 Å/3.4 Å/3.2 Å/2.9 Å by using a core mask. Reported resolutions are based on the gold-standard Fourier shell correlation (FSC) 0.143 criterion. Before visualization, all density maps were corrected for the modulation transfer function of the detector and sharpened by applying a negative B-factor that was estimated using automated procedures. Local resolution variations were estimated using RELION 3.0.

### Model building and refinement

The previously reported model (PDB: 5JP5) was used as the starting model and docked into the rCa_v_1.1-100S, rCa_v_1.1-10S, rCa_v_1.1, rCa_v_1.1-100R and rCa_v_1.1-100A maps in Chimera (*19*). The models were manually adjusted in COOT, followed by refinement against the corresponding maps by the phenix.real_space_refine program in PHENIX with secondary structure and geometry restraints. Statistics of 3D reconstruction and model refinement can be found in Table S1.

## Data and materials availability

Atomic coordinates and EM maps have been deposited in the PDB (http://www.rcsb.org) and EMDB (https://www.ebi.ac.uk/pdbe/emdb), respectively. For rCa_v_1.1 in complex with 100 μM, 10 μM (*S*)-(-)-Bay K8644, 1 μM (*S*)-(-)-Bay K8644 (apo in fact), 100 μM (*R*)-(+)-Bay K8644 and 100 μM amlodipine, the PDB codes are 7JPK, 7JPL, 7JPV, 7JPW, and 7JPX, respectively; EMDB codes are EMD-22414, EMD-22415, EMD-22424, EMD-22425, and EMD-22426, respectively.

## Acknowledgments

We thank the cryo-EM facility at Princeton Imaging and Analysis Center, which is partially supported by the Princeton Center for Complex Materials, a National Science Foundation (NSF)-MRSEC program (DMR-1420541). The work was supported by grant from NIH (5R01GM130762).

