## Supplementary Figures, Table, and Legends for "Structure Basis of Ca_v_1.1 Modulation by Dihydropyridine Compounds"

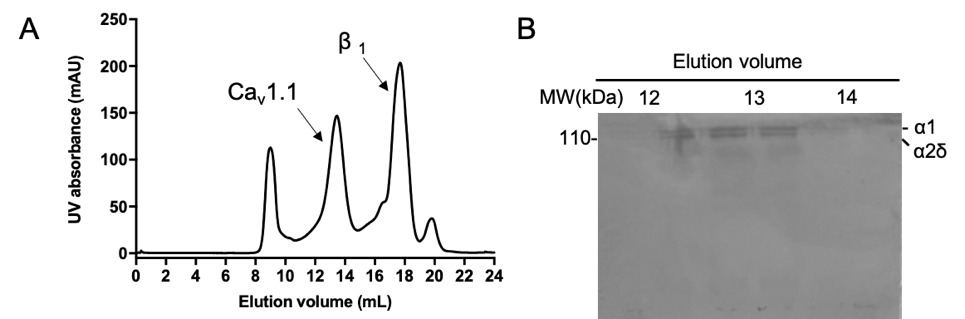

**Fig. S1 | Size-exclusion chromatography (SEC) purification of rCa_v_1.1.** (A, B) Representative size-exclusion chromatogram (A) and Coomassie blue-stained SDS-PAGE (B) for rCa_v_1.1 purified by glyco-diosgenin (GDN).

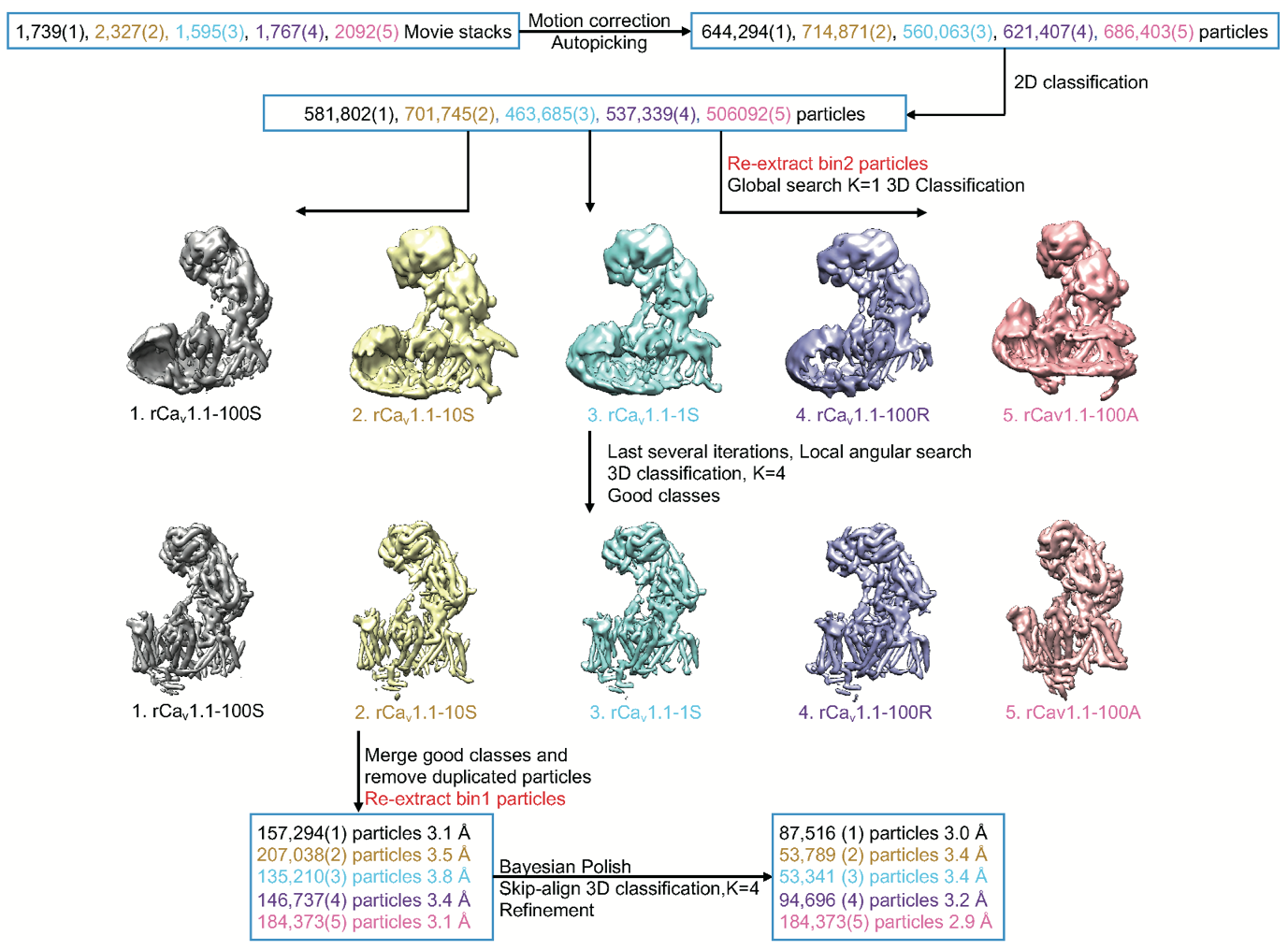

**Fig. S2 | Flowchart for EM Data Processing.** Details can be found in the image processing section in Methods. From left to right or from top to bottom: 1. rCa_v_1.1-100S (with 100 µM SBK); 2. rCa_v_1.1-10S (with 10 µM SBK); 3. rCa_v_1.1-1S (with 1 µM SBK); 4. rCa_v_1.1-100R (with 100 µM RBK); 5. rCa_v_1.1-100A (with 100 µM amlodipine).

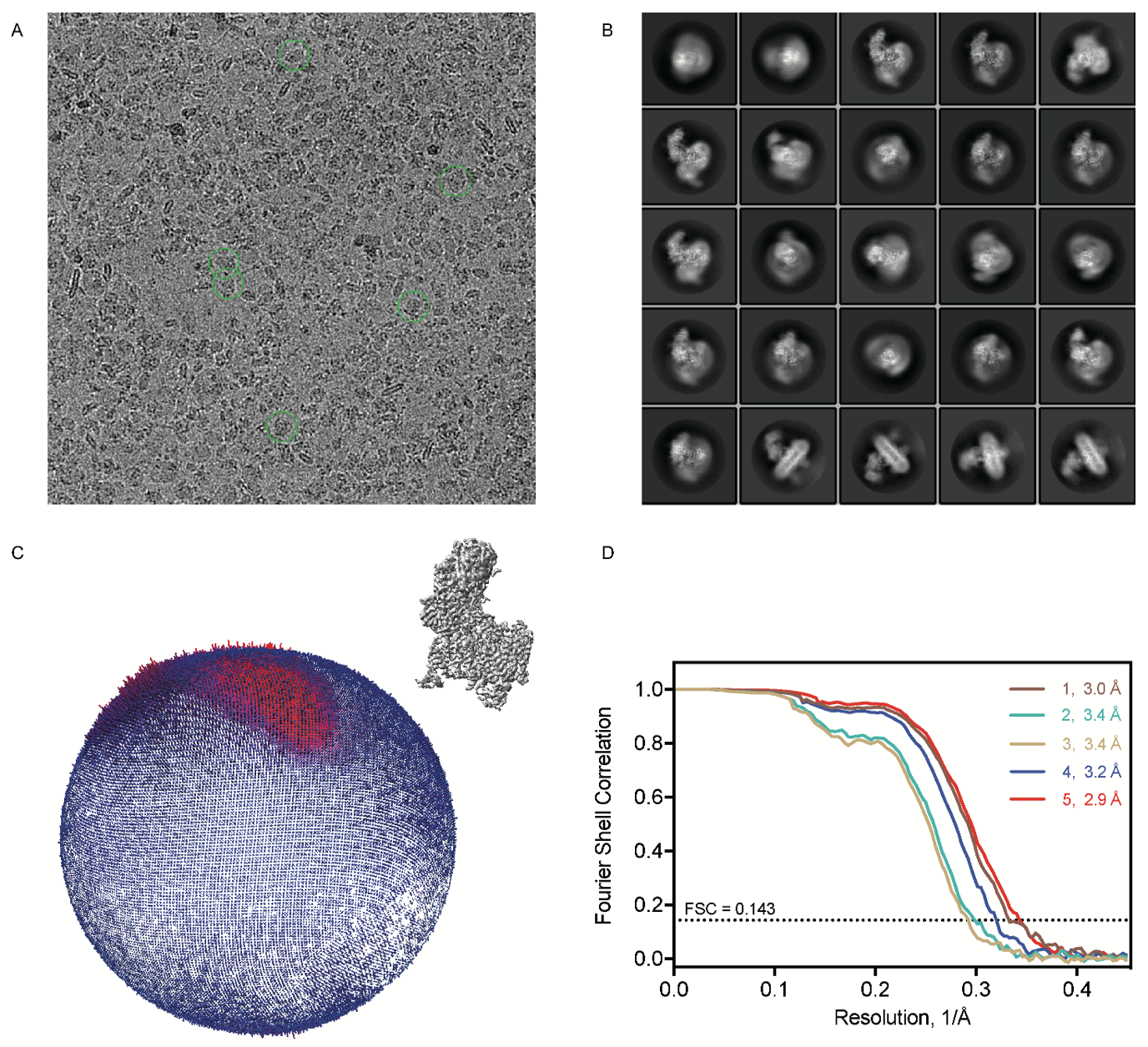

**Fig. S3 | Cryo-EM data analysis.** (A, B) Representative micrograph and 2D

class averages of rCa_v_1.1-100S in nanodiscs. Box size: 310 Å; circle mask: 260 Å. (C) Angular distribution of the particles of the final reconstruction of rCa_v_1.1-100S in nanodiscs (D) Gold-standard Fourier shell correlation (FSC) curves for the 3D EM reconstructions of 1, rCa_v_1.1-100S; 2, rCa_v_1.1-10S; 3, rCa_v_1.1-1S; 4, rCa_v_1.1-100R; 5, rCa_v_1.1-100A.

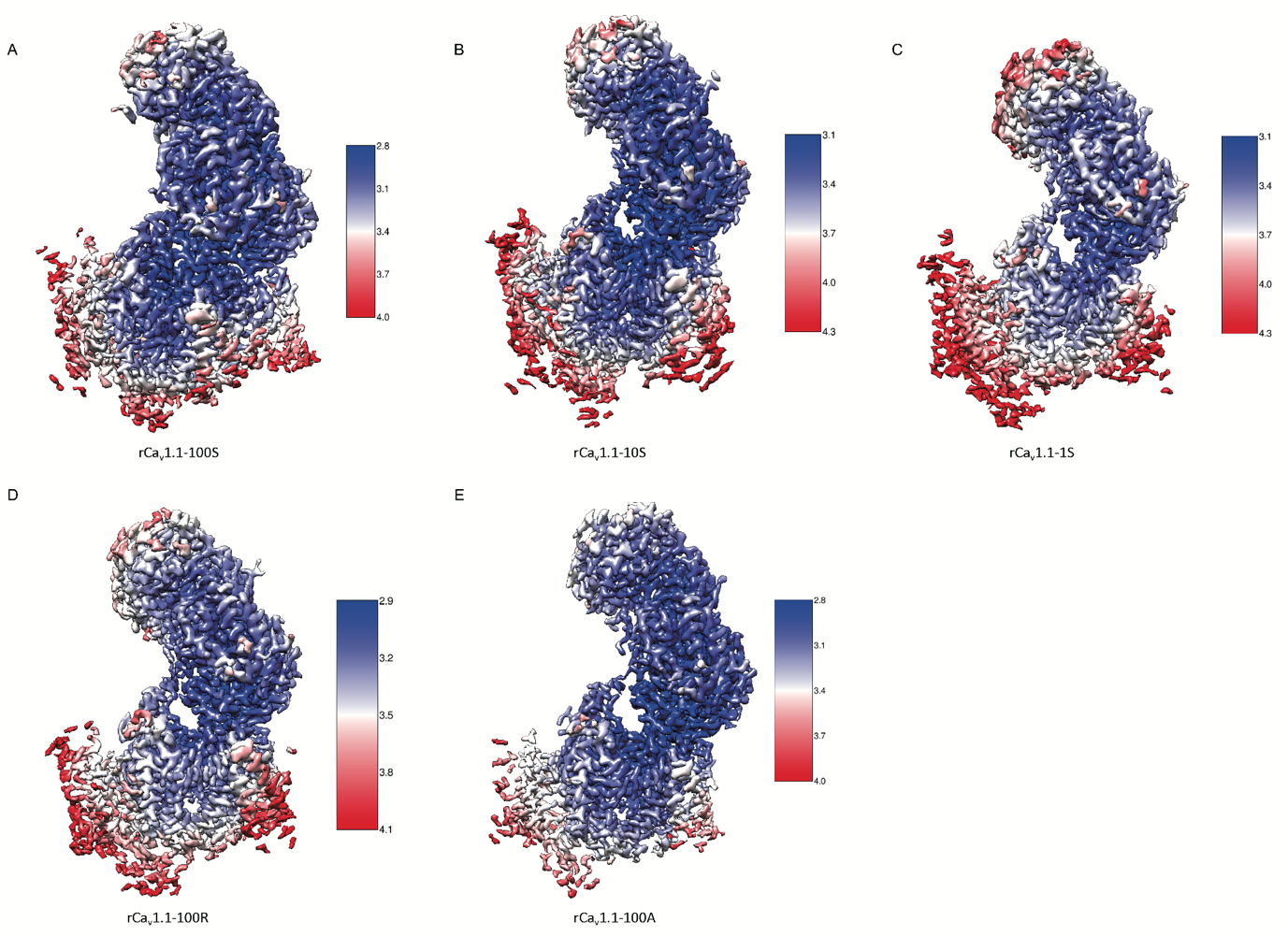

**Fig. S4 | Local resolution maps for rCa_v_1.1 reconstructions.** (A) rCa_v_1.1-100S, (B) rCa_v_1.1-10S (C) rCa_v_1.1-1S, (D) rCa_v_1.1-100R and (E) rCa_v_1.1-100A. The unit for the resolution scale bar is Å.

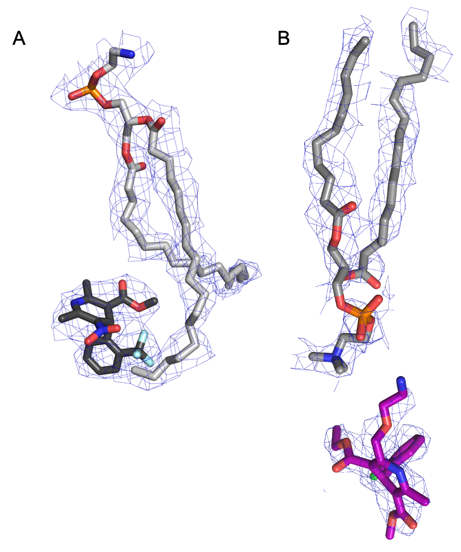

**Fig. S5 | Specific interactions between lipid and drugs.** (A) The density for a PD-surrounding lipid (PE, colored in grey) that interacts with SBK (colored in black) in rCa_v_1.1-100S. The maps, shown as blue mesh, are contoured at 2 σ for lipid and 3 σ for SBK. Please refer to Fig. 3E for the lipid. (B) The density for a traversing lipid (POPC, colored in grey) that interacts with amlodipine (colored in purple) within the pore domain of rCa_v_1.1-100A. The maps, shown as blue mesh, are contoured at 3 σ. These density figures were prepared in PyMol.

**Table S1 Summary of data collection and model statistics**

| **Dataset** | Ca_v_1.1-100S | Ca_v_1.1-10S | Ca_v_1.1-1S | Ca_v_1.1-100R | Ca_v_1.1-100A |
| --- | --- | --- | --- | --- | --- |
| EM equipment | Titan Krios (Thermo Fisher Scientific Inc.) | | | | |
| Voltage (kV) | 300 | | | | |
| Detector | Gatan K2 Summit | | | | |
| Energy filter | Gatan GIF Quantum, 20 eV slit | | | | |
| Pixel size (Å) | 1.114 | | | | |
| Electron dose (e^-^ /Å^2^) | 50 | | | | |
| Defocus range (μm) | 1.5 | | | | |
| # movie stacks | 1739 | 2327 | 1595 | 1767 | 2092 |
| Software | Relion 3.0 | | | | |
| Number of particles | 87,516 | 53,789 | 53,341 | 94,696 | 184373 |
| Symmetry | C1 | | | | |
| Resolution (Å) | 3.0 | 3.4 | 3.4 | 3.2 | 2.9 |
| Map sharpening B-factor (Å^2^) | -55 | -80 | -77 | -71 | -70 |
| Software | Phenix | | | | |
| Cell dimensions |  |  |  |  |  |
| a = b = c (Å) | 311.92 | 311.92 | 311.92 | 311.92 | 311.92 |
| α = β = γ (˚) | 90 | 90 | 90 | 90 | 90 |
| Model composition |  |  |  |  |  |
| Protein residues | 2257 | 2256 | 2227 | 2257 | 2258 |
| Ligands | 11 | 11 | 10 | 11 | 12 |
| R.m.s deviations |  |  |  |  |  |
| Bonds length (Å) | 0.005 | 0.008 | 0.004 | 0.014 | 0.012 |
| Bonds angle (˚) | 0.70 | 0.76 | 0.59 | 0.94 | 0.96 |
| Ramachandran plot statistics (%) |  | | | | |
| Preferred | 93.02 | 91.67 | 93.25 | 90.18 | 91.64 |
| Allowed | 6.84 | 8.29 | 6.57 | 9.73 | 8.22 |
| Outlier | 0.14 | 0.05 | 0.18 | 0.09 | 0.13 |
